# Cryo-TEM simulations of amorphous radiation-sensitive samples using multislice wave propagation

**DOI:** 10.1101/2021.02.19.431636

**Authors:** Benjamin A. Himes, Nikolaus Grigorieff

## Abstract

Image simulation plays a central role in the development and practice of high-resolution electron microscopy, including transmission electron microscopy of frozen-hydrated specimens (cryo-EM). Simulating images with contrast that matches the contrast observed in experimental images remains challenging, especially for amorphous samples. Current state-of-the-art simulators apply post hoc scaling to approximate empirical solvent contrast, attenuated image intensity due to specimen thickness, and amplitude contrast. This practice fails for images that require spatially variable scaling, *e.g.*, simulations of a crowded or cellular environment. Modeling both the signal and the noise accurately is necessary to simulate images of biological specimens with contrast that is correct on an absolute scale. The “Frozen-Plasmon” method is introduced which explicitly models spatially variable inelastic scattering processes in cryo-EM specimens. This approach produces amplitude contrast that depends on the atomic composition of the specimen, reproduces the total inelastic mean free path as observed experimentally, and allows for the incorporation of radiation damage in the simulation. These improvements are quantified using the matched-filter concept to compare simulation and experiment. The Frozen-Plasmon method, in combination with a new mathematical formulation for accurately sampling the tabulated atomic scattering potentials onto a Cartesian grid, is implemented in the open-source software package *cis*TEM.

## Introduction

The power (variance) of the noise in cryo-EM images outweighs the power of the signal, often by a factor of 20 or more. The dominant source of noise in cryo-EM is “shot” noise, arising from the stochastic nature of detecting an electron at a given location and time due to low-dose imaging conditions. A detailed analysis by Baxter et al [1] demonstrated the need to also consider structural noise, defined as any contrast arising from sources other than the final object of interest: carbon film, crystalline ice, radiation damaged particles, unwanted macromolecular conformers, the supporting amorphous ice etc. Unlike shot noise, structural noise is affected by objective lens aberrations, which give rise to the contrast transfer function (CTF). Baxter et al. modelled both the structural noise and the shot noise as additive white Gaussian noise, which fails to capture the artifacts and challenges that are commonly encountered during image processing, as previously demonstrated by Scheres et al [2].

An improvement in how the structural noise is simulated, particularly that arising from the supporting amorphous ice, can be found in TEM simulator [3] and inSilicoTEM [4]. They implement multislice wave propagation as described originally by Cowley and Moodie [5], resulting in noise that is affected by the CTF. The result of a multislice simulation is a probability distribution defined by the squared complex modulus of the electron wave function at the detector *ψ_d_(x, y)*. The simulated image is then formed by drawing from a Poisson distribution unique to every pixel while incorporating the influence of the detector quantum efficiency (DQE).

Most of the information transferred from the specimen to the image in high-resolution cryo-EM is captured in phase contrast arising from interference between elastically scattered and non-scattered electrons. Amplitude contrast is also present due to electrons scattered outside the objective lens aperture, and loss of electrons from the elastic image due to inelastic scattering. The latter source of amplitude contrast is enhanced using an energy filter [6]. Amplitude contrast cannot be explained by linear image formation theory [7] and is accounted for post hoc via a phase shift term added to the CTF applied to the simulated image [8]. This treatment is also common practice in solving the inverse problem of image reconstruction, which seeks to answer the question “*what is the probability of the model given the observed data*.” However, in forward modeling, which asks “*what is the probability of observing some data given a particular model*,” it would be desirable to account for the fact that amplitude losses depend both on atom type, and on local mass thickness. For example, heavy atoms like gold scatter more electrons outside the objective lens aperture than light atoms. On the other hand, light atoms produce more amplitude losses in energy filtered images than heavy atoms due to their higher ratio of inelastic to elastic scattering [9][10].

The multislice formalism is essential for thick specimens where the projection approximation fails, as it incorporates important effects like multiple scattering of electrons and the curvature of the Ewald Sphere. Increasingly thick samples are also less transparent to electrons, and all simulators we are aware of apply an implicit “energy filter” to remove inelastically scattered electrons from the final image. To account for inelastic losses, a single thickness parameter is used to attenuate the image intensity according to

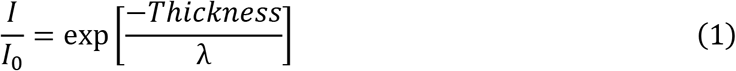

where *I*_0_ is the unattenuated image intensity, and λ is the inelastic mean free path for single scattering - the average distance an electron passes through the specimen before scattered inelastically at least once. It is clear that these single filters cannot work for specimens with variable mass thickness, *e.g.*, at the edge of a cell, for variable atomic composition, *e.g.*, the increased phosphorus concentration in the nucleus, and even for many single particle samples [11].

Even with a limited subset of atomic species, to which we will constrain the following discussion, there are two very different environments that need to be simulated - the molecule and the solvent. We will refer to how well the molecule stands out from the solvent as “solvent signal-to-noise-ratio or *SNR_solvent_*” as quantified by Yonekura et al where *Ī* is the mean image intensity and *σ_solvent_* is the standard deviation in the solvent region.

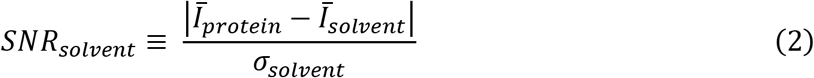

Typically, the solvent is added on top of the simulated molecules in projection, with a single value given by the mean inner potential for aqueous water. This approach, which we will refer to as “the continuum model”, is equivalent to using an infinite time average of a collection of moving water atoms. One shortcoming of the continuum model is the failure to account for the hydration radius of a molecule, which should be zero inside a particle, higher than the bulk solvent immediately outside the particle envelope and gradually falling off with distance [12]. Ignoring the fact that molecules displace the solvent has been shown to produce *SNR_solvent_* that fails even visual inspection at exposures of 100 e^−^/Å^2^ [4].

We now know that the infinite time average used in the continuum model does not adequately describe reality; even though the solvent is frozen low-density amorphous ice (LDA), it is not static during the imaging process. McMullan and Henderson quantified the motion of water molecules in LDA during imaging, estimating a RMSD of 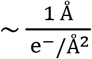 [13]. Importantly, this motion results in a blurring of the solvent over time, which can be thought of as low pass filtering, and so *σ_solvent_* decreases with increasing exposure. The net result is that *SNR_solvent_* is a function of the total exposure in an image, gradually increasing as the solvent becomes more blurred. Of note, the increase of *SNR_solvent_* with exposure is further amplified in experimental images by mass loss, which also decreases *σ_solvent_* and increases the numerator in Equation 2 by reducing *Ī_solvent_*. A more sophisticated version of our solvent model may implement this mass loss in future work.

While *SNR_solvent_* is useful for its simplicity, a more detailed analysis requires another metric to quantify how well the macromolecules we simulate match experimental images. For this, we propose using the matched filter, which is the statistically optimal realization of a cross-correlation detector. With image statistics characteristic of cryo-EM data, the output of the matched filter can be simply defined as the ratio of the cross-correlation coefficient *(CCC)* to the standard deviation of the *CCC* when only noise is present (*σ_n_*) [14] including any sources of structural noise as defined above.

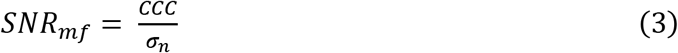

The upper bound on the *SNR_mf_* is given by the ratio of the power of the input signal to the power of the noise in the image [15]. This means, for example, that a larger molecule will generally have a higher *SNR_mf_*, while any disagreement between the signal in the image and the simulated template reduces the *SNR_mf_* from this maximal value. As such, the relative accuracy to which the simulated molecular density matches experimental data can be determined by searching images using a matched filter. To evaluate Equation 3, we use the cross-correlation tools and relevant preprocessing as available in *cis*TEM [16][17].

To understand why the continuum model fails to produce accurate *SNR_solvent_* fails to produce accurate it is instructive to consider the source of the motion of the solvent upon exposure to the imaging electrons. Along with radiation damage to the molecule of interest, sample motion is the result of energy being transferred to the specimen via inelastic scattering. For the samples in which we are interested, inelastic scattering is generally attributed to plasmons, i.e., collective excitation of valence electrons by the electric field of the imaging electrons. However, the extent to which these are bulk plasmons, which are strongly delocalized, or more localized single-electron excitations remains unclear [18]. Independent of the exact form of the plasmons, their net effect is an alteration of the system’s Hamiltonian during imaging, such that product of a traditional multislice simulation, *ψ_d_(x, y)*, is no longer valid. Just as the original multislice method introduced a division of the specimen potential into thin spatial slices to ensure the small angle approximation is valid, we suggest dividing the simulated exposure into small temporal slices, where the specimen does not change too much. While we refer to time here, what is most practical from the microscopist’s point of view is exposure measured in e^−^/Å^2^. Therefore, the time step in our simulator is specified as the desired exposure per movie frame. Exposure rate dependent phenomena like detector DQE and beam coherence are parameterized by an exposure rate with the exposure time implicitly set by the software according to the user supplied exposure-per-frame divided by the exposure rate. The DQE parameterization is based on the results of Ruskin et al [19] while the coherence is parameterized based on the beam brightness of an Xfeg as given in the Titan Condenser Manual [20].

## Theory

There are three main components in modeling the image formation process in high-resolution transmission electron microscopy (HRTEM) of which cryo-EM is a subset:

1. The relativistic electron wave function and its modulation by the sample.
2. The exposure-dependent Coulomb potential of the specimen.
3. The microscope, including apertures, detector, lens optics and aberrations.

In this work, we are concerned primarily with how the Coulomb potential changes due to energy being deposited in the specimen during imaging and will provide only a brief summary of the other two components. The interested reader is referred to, in increasing order of completeness, the treatments by Vulović et al [21], Kirkland [22], Reimer and Kohl [23] Hawkes and Kasper [24].

Unlike photons, electrons have a spin quantum number and so their interaction with matter is governed by solutions to the Dirac equation. Given reasonable approximations [24], a relativistically corrected version of the Schrödinger wave equation, called the Klein-Gordon equation is used in practice.

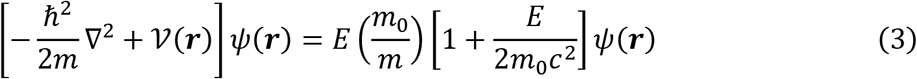

Analytical solutions to this equation are intractable for all but the simplest systems, so we turn to multislice wave propagation [25], which produces an approximate numerical solution to this equation. The first step in a multislice simulation is the calculation of the specimen’s Coulomb potential *ν(**r**; t)*. The time dependence will be omitted assuming a quasi-stationary solution for an exposure to a single electron. The potential is divided into thin slices along the imaging axis, which can be approximated by two-dimensional scattering potentials through which the electron wave function is sequentially propagated. This subdivision ensures the potential varies slowly in the direction of the electron wave propagation, such that the small angle approximation remains valid and scattered spherical wave fronts may be approximated locally by a parabola (Fresnel diffraction.) In the limit of infinitely thin slices, this results in an exact numerical solution to the Klein-Gordon equation [26].

Multislice simulations can model both elastic and inelastic scattering processes, provided that the respective Coulomb potentials can be calculated. In analogy to the optical potential in light microscopy, inelastic scattering is incorporated into the wave theory via a complex term in the specimen potential as introduced by J.C. Slater in 1937 [27].

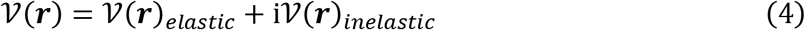

The isolated atom superposition approximation states that the specimen potential *ν(**r**)* may be represented as the sum of the individual atomic potentials *φ(**r**)_i_*. We introduce a scaling factor β to compensate for the contribution of bonds among those atoms to maintain the correct total scattering cross section:

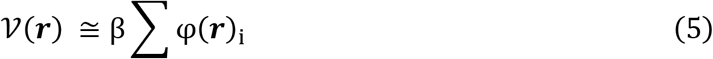

The elastic atomic potential can be calculated using relativistic Hartree-Fock wave functions [28]. The solutions for isolated atoms, having isotropic distributions, are commonly parameterized by a sum of four or five Gaussian functions [29]. Typically, the potential is recorded indirectly as these fits are tabulated as elastic electron scattering factors, defined as the Fourier transform of the elastic potential [30].

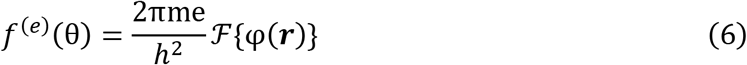

The numerator describes the product of the electron charge and relativistic mass, *h* is the Planck constant, and 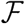 denotes the Fourier transform operator. An important relation we will return to later equates the spectral distribution of the scattering factor to the differential scattering cross section - the probability of an electron being scattered through a solid angle Ω:

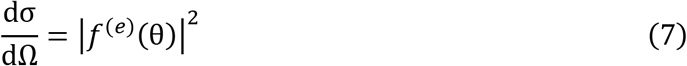

While *ν(**r**)_inelastic_* is straightforward to calculate from first principles, *ν(**r**)_inelastic_* is more problematic given the varied mechanisms with which an incident electron may transfer energy to the specimen: ionization, excitation, dissociative attachment, vibrational and rotational excitations, bremsstrahlung, etc. [31]. One example where *ν(**r**)_inelastic_* is well defined is for radiation-insensitive crystalline specimens, where thermal diffuse scattering (TDS) caused by phonon excitation is the primary contributor to the complex potential [29]. One model to calculate the TDS potential treats the time average atomic displacement through Debye-Waller factors and improves the accuracy of dynamic RHEED calculations [32]. This time average approach is analogous to how the solvent calculation, specimen motion, radiation damage and alignment errors are accounted for in HRTEM simulations of biological specimens by B-factors, which are related to Debye-Waller factors by a factor of 4.

While this time-averaged approach preserves the total intensity of the projected interaction potential [33], it is well known that the image contrast produced in this way is systematically wrong, often by a factor of three or more. The error, known as the Stobbs factor [34], becomes worse with an increasing strength of the electron specimen interaction, which in turn, depends on the average mass thickness in the specimen. Stobbs et al. proposed two likely causes for the observed contrast mismatch between simulation and observation: a) existing simulators do not account properly for radiation damage to the specimen, and/or b) they fail to model the inelastic scattering with sufficient accuracy. As recently shown empirically, these are related phenomena [35].

Van Dyck et al. demonstrated that the Stobbs factor could be largely corrected for by using the “frozen phonon” method [36][37]. The approach is conceptually simple: A series of simulations are carried out where each atom is displaced randomly based on empirical TDS values. The intensities in the image plane as calculated from these individual simulations are then averaged together. Here we propose a similar idea, applied to radiation-sensitive frozen-hydrated specimens, where plasmons are the primary form of inelastic scattering. The “frozen plasmon method” presents several computational and theoretical challenges:

1. The number of solvent atoms (O(10^9^)) greatly outweighs those of the macromolecules we wish to simulate (O(10^5^)), requiring careful algorithmic design to make the computations tractable.
2. The solvent and macromolecules have very different elastic and inelastic total scattering cross-sections, as well as different average mass densities (~0.94 g/cm^3^ for low-density amorphous ice and ~1.38 g/cm^3^ on average for proteins) [38]. This means that the amplitude contrast and inelastic losses cannot be applied ad hoc to the final simulated image and must be considered on a per-atom basis.
3. The preceding points also place a requirement on the accuracy of the calculation of each atomic scattering potential, which can no longer simply be rescaled and so must be correct from the start.
4. The scattering factor for plasmons in low-density amorphous ice is needed to achieve the appropriate contrast, which depends on the appropriate spectral distribution. To obtain an expression ϕ^(inelastic)^*(**r**)_i_*, we start from the double differential scattering cross-section for plasmons [39]. The essential form is Lorentzian

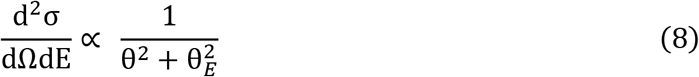

with the angular dependence θ and the energy dependence captured in the characteristic angle

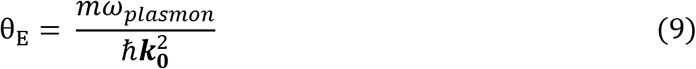

*ℏω_plasmon_* is the energy loss of the plasmon, and ***k*_0_** the incident electron’s wave vector. To calculate a scattering factor for plasmons, we first form an empirical probability distribution for plasmons arising from singly scattered electrons in amorphous ice, derived from EELS published by Du and Jacobsen [40]. We then numerically integrate Equation 8 over energies in the low-loss spectrum (7.5 - 100 eV) for each angle. We then combine this spectrum with empirical measurements of the ratio of inelastic to elastic total scattering cross sections, which are inversely proportional to atomic molecular weight [10]. As we calculate the elastic potential during simulation, we separately accumulate an inelastic potential scaled per atom by these total probabilities. During wave function propagation, this inelastic potential is given the correct Lorentzian form, taken to be the square root of the values above.

Plasmons scatter strongly at low angles and are generally referred to as being delocalized. This is reflected in Figure 1 where the inelastic scattering factor we derived for plasmons is compared to the elastic scattering factor for a glutamine molecule. While plasmon scattering dominates at low resolution compared to elastic scattering, it still contributes significantly at high angles as can be seen by the red hash marks in Figure 1 that demarcate bins of 20% total inelastic scattering probability. The precise nature of inelastic scattering in amorphous materials is not well understood, such that the relationship between this high-resolution information and the underlying specimen structure is not defined.

**Figure 1.**
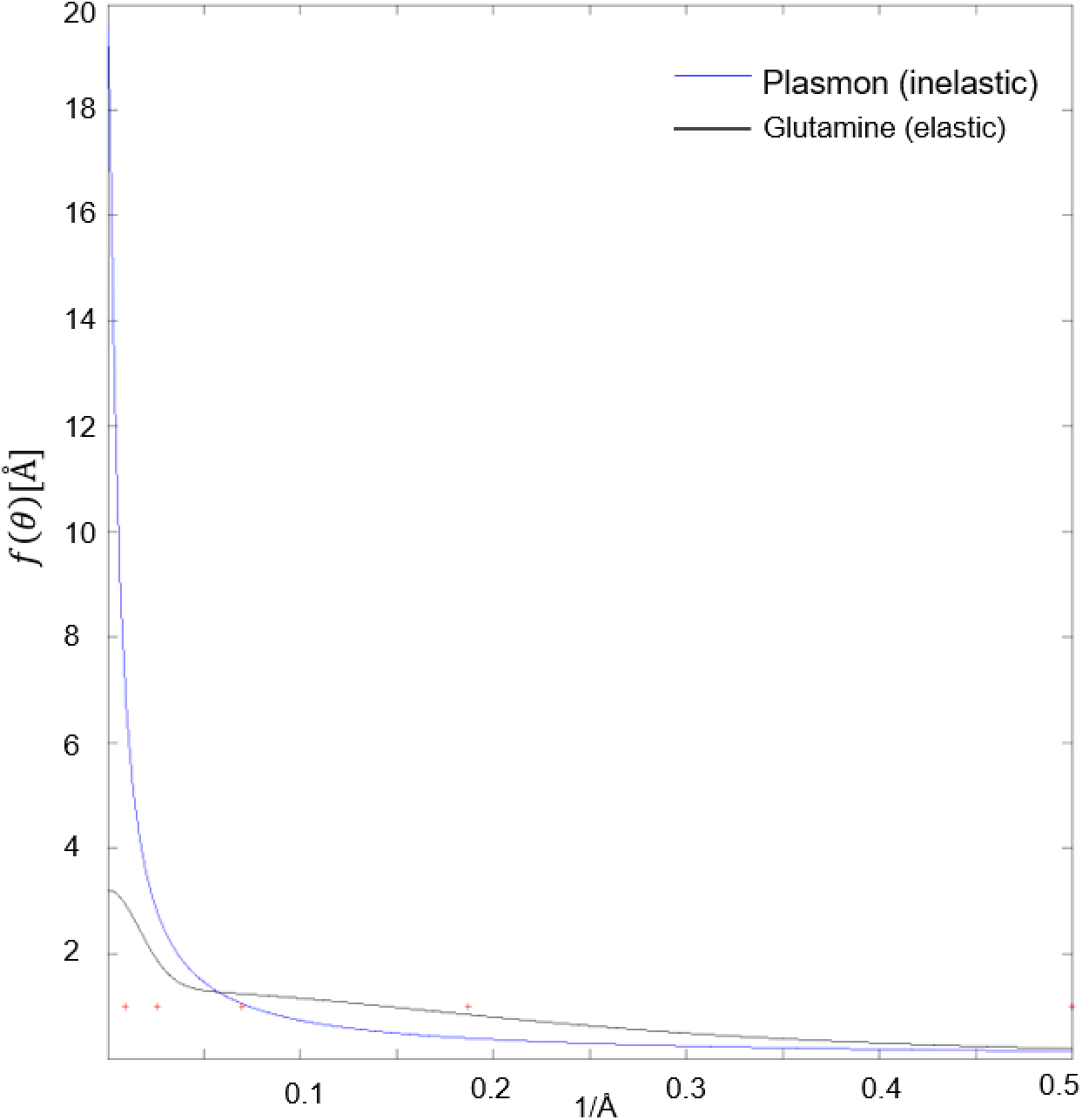
Comparison of elastic scattering factor for a glutamine molecule (black line) vs. the inelastic scattering factor for a plasmon pseudo-particle (blue line.) Red markers indicate the right edge of bins comprising 20% total scattering probability for the inelastic Plasmon scattering factor as calculated from the squared norm of the scattering factor.

## Results

### Accurate representation of molecular density

For isolated neutral atoms, the scattering potential, defined as the Fourier transform of the parameterized scattering factors, can be written as

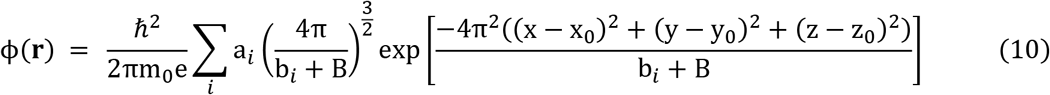

This atomic potential is sharply peaked in real space, requiring a high sampling rate when discretizing in order to maintain the total projected potential. This high sampling rate effectively produces a numerical integration of Equation 10. To allow for coarser sampling, and hence improve the efficiency of our simulator, we analytically integrate the expression from Equation 10.

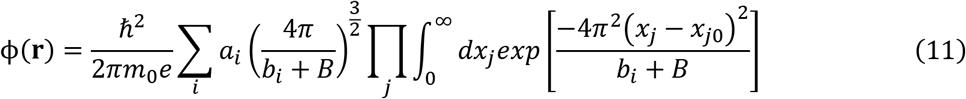

resulting in

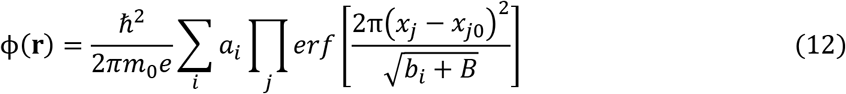

While the potential in each voxel is marginally more complex to calculate (to evaluate the limits of integration, the error function must be evaluated six times per voxel, compared to a single exponential) this is more than compensated by the reduced number of voxels needed. For example, simulating at 0.5 Å voxel pitch is 125x less computationally expensive than simulating at 0.1 Å voxel pitch. While the voxel pitch is the same in the z-dimension, the slab thickness is a free parameter which also affects computational efficiency. A simple test to determine the maximum allowable thickness, as suggested by Kirkland [22], is to search for the point where the results of the simulation become dependent on slab thickness. Our simulations begin to show a dependence on slab thickness around 7 Å (data not shown) and, therefore, we typically use 5 Å. Even more important than computational speed, using Equation 12 in our simulations also means the sampled potential still has the correct magnitude and is not simply proportional to the continuous potential, as discussed in the following section.

### Compensating for the isolated atom superposition approximation

While the integral formulation of the scattering potential preserves the calculated potential of all the individual atomic contributions, there is still a systematic underestimation of the scattering potential due to bonding interactions. This is generally estimated to be between 5-10% of the total potential [41], and ignoring this difference is referred to as the isolated atom superposition approximation. Given that we want to obtain images that are quantitative on an absolute scale, we sought to measure and calibrate this error. To approximate the redistribution of the scattering potential due to bonding in a biological specimen, we use the available data for amorphous carbon comparing to results from electron holography as follows. The average phase shift in a material depends on the mean inner potential of the material (*V*_0_), the thickness (*t*), and an interaction constant *C_E_* [23].

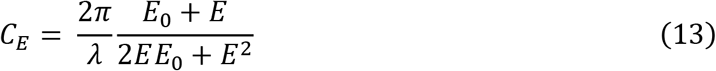

Additionally, surface boundary effects are also known to be important in cryo-EM imaging, so we compared our calculated phase shift (*δφ*) to empirical results obtained using electron holography, which measures both the mean inner potential of carbon (*V*_0_ = 9.04*eV for* 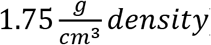) and an additional thickness-independent surface-induction phase shift *φ_add_* (0.497 radians)[42].

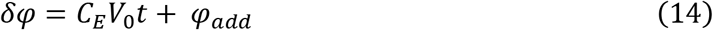

Considering the principle of a Zernike phase plate, we simulated an amorphous carbon sheet that should produce a phase shift of *π*/2 radians (Figure 2A) with a density of 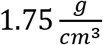 and 348.6 Å thickness per Equation 14. Our simulation suggested that the average phase shift is ~3.8% too small. To correct for this error, we introduce a constant scaling factor (*β*) of 1.038 to the isolated atomic potentials. The simulated phase plate also serves as a sanity check that the calculation of the elastic scattering potential is consistent across different pixel sizes (Figure 2B).

**Figure 2.**
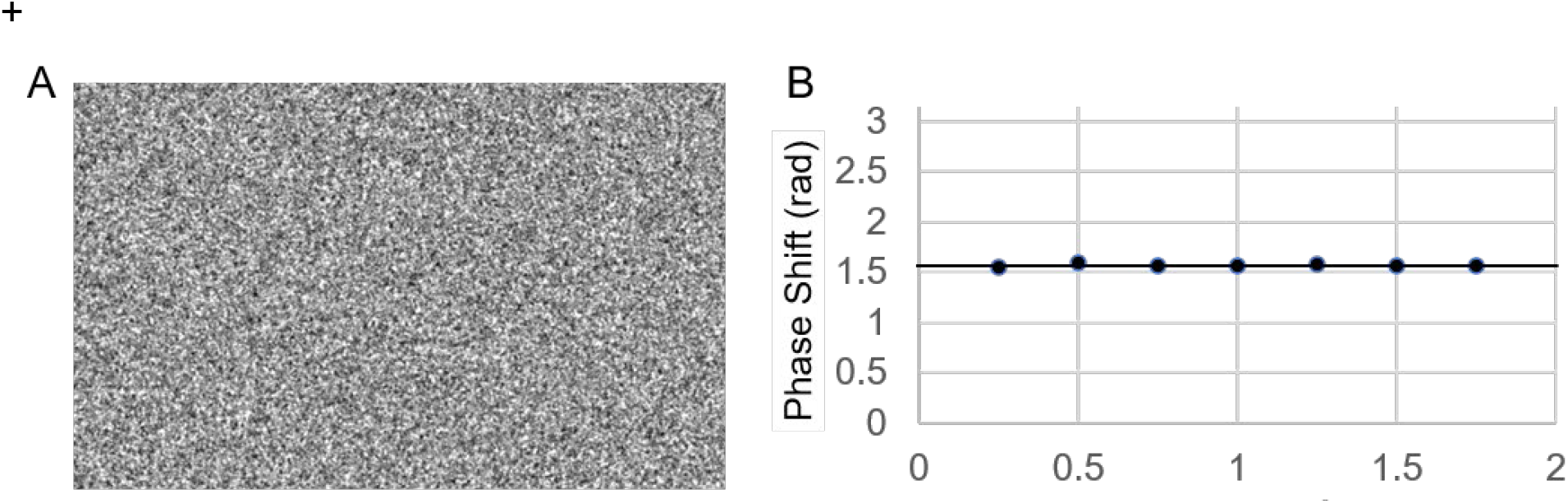
A) “phase plate” simulated from amorphous layer of carbon atoms 348.6Å thick with a density of 1.75 g/cm^3^ B) Mean phase shift for the simulated phase plate as a function of pixel sampling rate during simulation. Black line plotted at *π*/2 radians for visual reference.

+

### Modeling the solvent envelope

In the previous section, we demonstrated that we could accurately calculate the contrast for a collection of randomly distributed atoms of a given density. Next, we show that in order to accurately compare a simulated protein density to experimental data, we must also consider the solvent displaced by the protein. This creates a low-resolution “hole” that impacts subsequent analysis as discussed in detail by Zhang and Sigworth [12]. We incorporate their hydration radius model into the simulator by tracking the smallest distance to any non-solvent molecule and weighting any nearby solvent with a probability distribution defined by normalizing Equation 1 from their paper. (We note that the parameter “r3” in table 1 of Zhang and Sigworth should be ~3.0, not 1.7, personal communication.)

When simulating isolated molecules to use for comparison to experimental images, we weight the average water potential by this probability distribution, with an exponential decay beyond 4 Å. This exponential decay is added because our knowledge of the sample rapidly decays to zero beyond the particle of interest. This produces an effect similar to the ad hoc model suggested previously by Henderson and McMullan [43]. When simulating images, the probability distribution is applied to individual pseudo-water molecules as described next.

### Modeling radiation damage

As soon as the electron beam is “switched on” the sample begins to accumulate radiation damage. This has long been known to be *the* limiting factor in cryo-EM [44] and an analytical function describing the effects of radiation damage as a Fourier space filter - *ξ(**q**)* - was described by Grant and Grigorieff [45]. Since radiation damage is specimen dependent, the analytical model of Grant and Grigorieff will only strictly apply to rotavirus VP6 capsid protein, and not to nucleic acids, for example. Alternatively, radiation damage combined with other errors, for example uncorrected motion blur, may be fit using exposure-dependent B-factors [46][47]. The latter approach has the advantage that it does not try to separate the blurring due to radiation damage from other sources of blurring, which can be difficult in practice.

To quantify the accuracy in modeling radiation damage using the analytical model, *ξ(**q**)*, we employ the matched filter concept which is sensitive to the spectral distribution of the template when the noise does not have a flat power spectrum. Using the rotavirus DLP images originally used by Grant & Grigorieff to derive their radiation damage model, we investigated several sources of such noise including the water shell, as described in the next section, the detector modulation transfer function (MTF), CTF coherence envelope, residual intra-frame motion, atomic modeling uncertainty and defocus uncertainty.

By looking at a single early frame from each of 18 movies, each with ~1.5 *e*^−^Å^2^ total exposure (0.77 *e*^−^Å^2^ intra frame exposure), we could quantify how well we modelled these sources of noise, while assuming no significant radiation damage. In Figure 3A we show the projected scattering potential of our simulated DLP (PDB 3gzu) along with the average *SNR_mf_* from 180 DLPs overlaid. Each source of noise is included cumulatively, showing the relative improvement in detection as the model becomes increasingly more detailed. Applying additional positive or negative B-factors only made the score worse. To illustrate the effect of each of these modifications on the template’s spectral distribution, we also plot the ratio of the rotationally averaged power spectra of each simulation to the previous one. A representative image for frame 2 is shown in Figure 3B, while an average of frames 2-91 is shown in Figure 3C.

**Figure 3.**
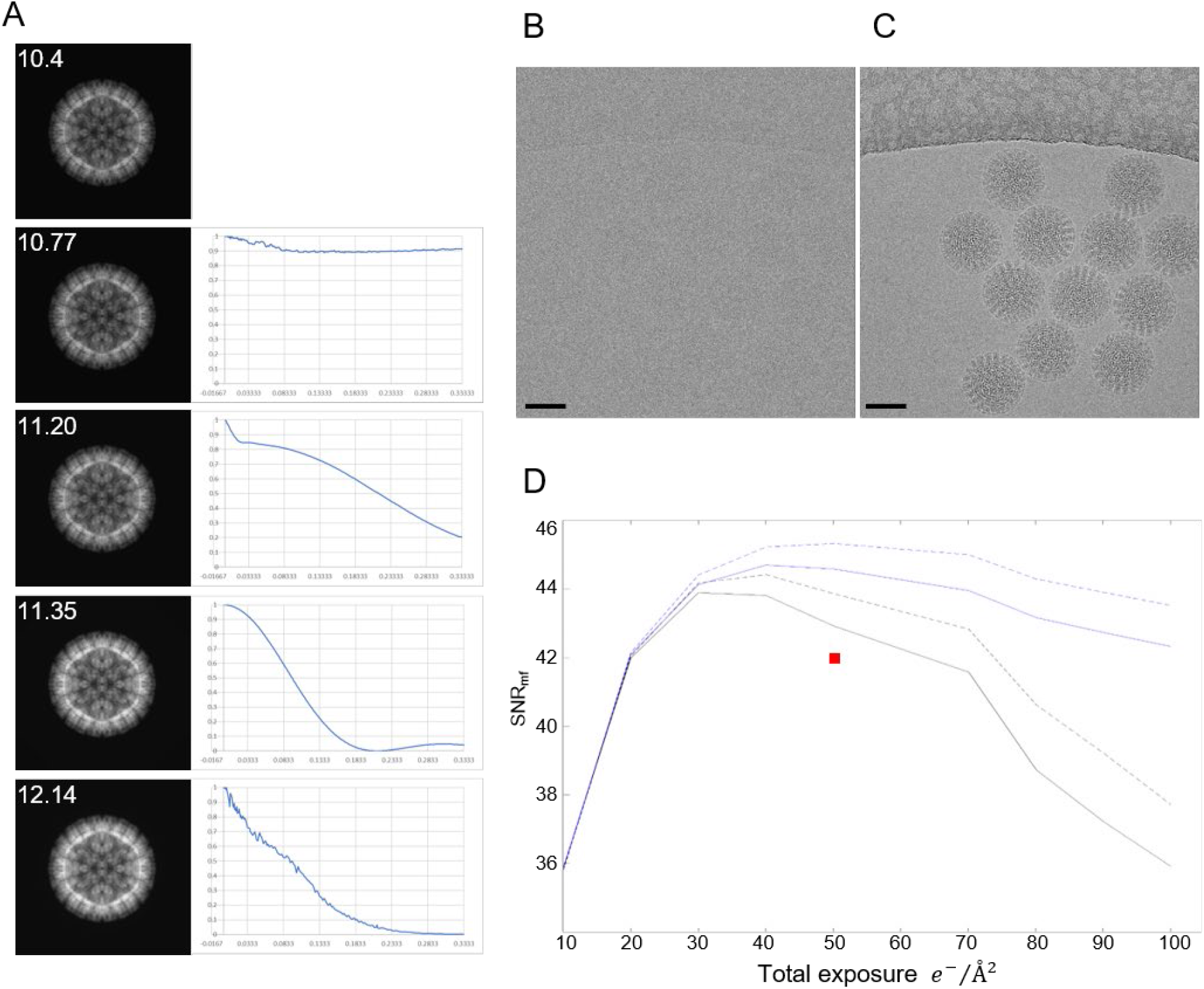
A) Left column, projections of simulated rotavirus DLPs from PDB-3gzu with each of the blurring process applied cumulatively. Average peak *SNR_mf_* from 180 DLPs over 18 images overlaid in white. Right column, the shape of the individual envelopes produced by the respective blurring as calculated by dividing the rotationally averaged power spectrum by the preceding image. B) Example image from frame 2 and C) frames 2-91 of DLPs from the data set used in this analysis, kindly shared by Dr. Tim Grant. D) Average SNR values as a function of total exposure, frames averaged from 2-N, where N is on the total exposure on the horizontal axis. Solid black line, no filtering. Dashed black line, image summed from exposure weighted movie frames. Solid blue line, reference simulated with cumulative exposure filter as would normally be applied to a movie. Dashed blue line, both image and reference exposure filtered. Red square is the maximum SNR obtained by using a single B-factor to represent all the envelopes in (A). Scale bars 500 Å.

To assess the radiation damage model, we averaged movie frames 2-N such that the accumulated exposure ranged from 10 – 100 *e*^−^Å^2^ either with, or without, exposure filtering applied to the images. We then measured the *SNR_mf_* of the DLPs in these two sets of images using two sets of references calculated with, and without, exposure filtering applied during simulation, and plotted the results in Figure 3D. We found the largest increase in *SNR_mf_* using a total exposure of 50 *e*^−^Å^2^ when applying the exposure filter to both the image and during simulation of the reference. To account for blurring due to residual intra-frame specimen motion in this experiment, the exposure filter was additionally modified to include a damping envelop which, for uniform motion, is trivial to show is a sinc function. Less trivial is determining the intra-frame specimen motion (***d**_i_*) for which we only know a lower bound, estimated as the average of the displacements between the two neighboring frames, and applied as a sinc modulation along with the exposure filter (if included) for that frame.

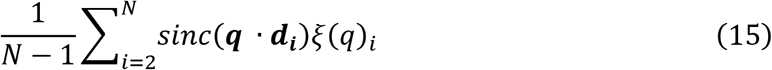

Taken together, these results suggest that other modifications to the template that result in a better match to the experimental data can further improve the *SNR_mf_*; for example, some amino acid side chains are affected more strongly by radiation damage than others, *e.g.*, aspartate and the disulfide bond of cystine. These details could be incorporated into a new atom specific damage model in future work.

### Accurate representation of solvent noise

Because the number of water molecules is very large, we elected to calculate a coarse-grained model for water, where each water molecule is represented as a single, isotropic scattering center. We based the scattering factor for our pseudo-waters on the elastic scattering factor tabulated for oxygen, but scaled by the ratio of the total elastic scattering cross-section of oxygen:water, which we know from experiment [31]. These pseudo-molecules are seeded randomly at the proper density for low density amorphous ice (~0.94g/cm^3^). A movie is then simulated, where each time step (movie frame) is defined by a user-specified exposure and the specimen is held constant within that time. The simulated probability density for the constant potential is shown in Figure 4A while our coarse-grained all-atom model is shown in Figure 4B. The average intensity in the solvent region is the same in both images.

**Figure 4.**
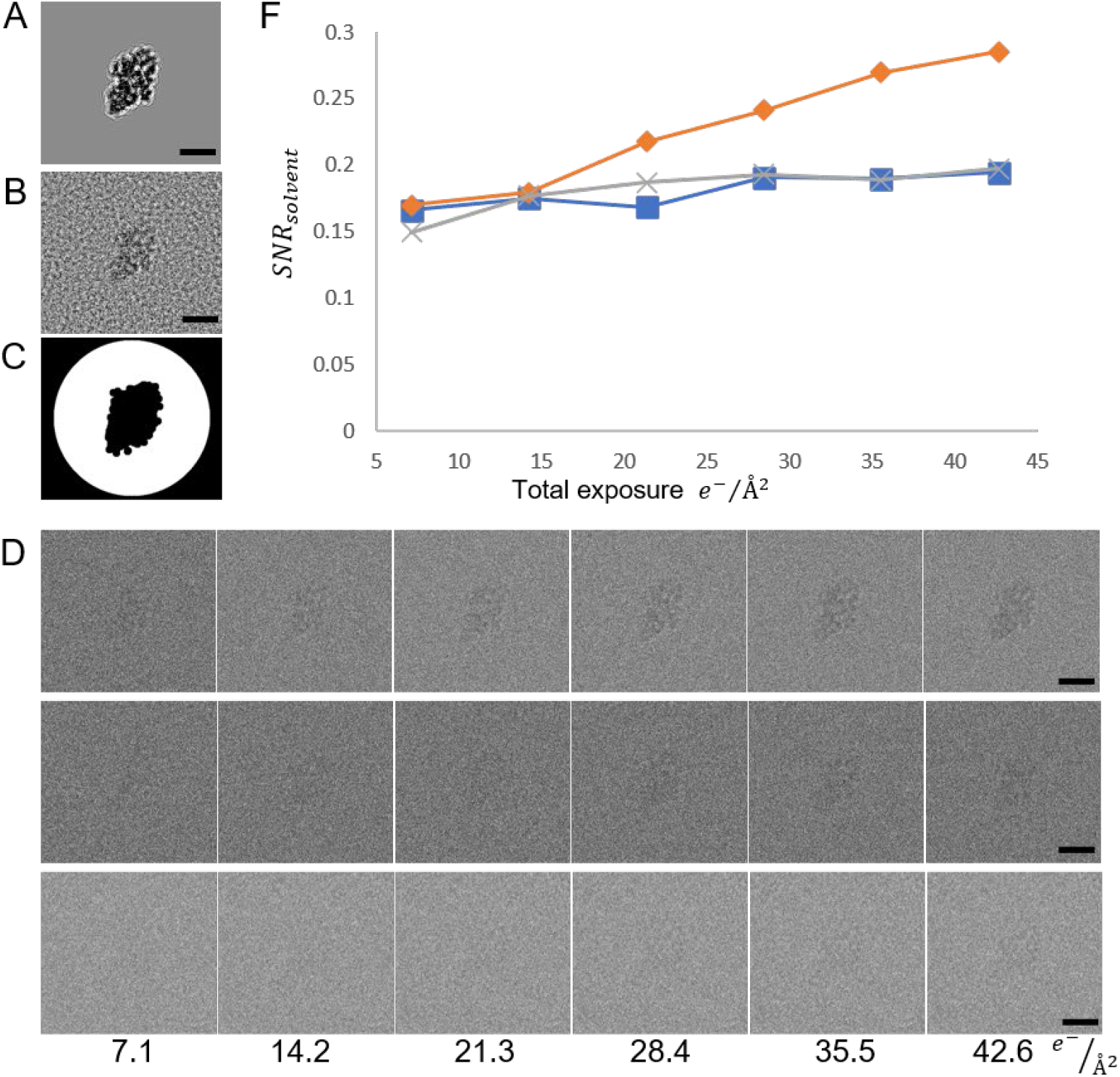
Comparing the continuum and coarse-grained all-atom solvent model to experimental data. A) │Ψ*_detector_*│^2^ with a constant potential added for the solvent. B) │Ψ*_detector_*│^2^ with coarse-grained all-atom water model. The average intensity in A and B in the solvent region is identical within numerical precision. C) Mask used in calculating *SNR_solvent_* (equation 2 main text) where the white region was used for the solvent, and the central black region for the protein. D) Plot of *SNR_solvent_* as a function of accumulated exposure for experimental data (Blue line, square marker) coarse-grained all-atom solvent model {grey, x-marker) and constant solvent potential (orange, diamond marker). E) Images used in calculating (D) with the same color/marker scheme, and total exposure indicated along the bottom. Experimental data taken from EMPIAR-10061, beta-galactosidase. Scale bars 100 Å

In Figure 4D we show selected time points from a movie simulated using the continuous solvent potential model (top row), the coarse-grained all-atom model (middle row), and experimental data in the bottom row (EMPIAR-10061 [48]). As can be seen visually, the *SNR_solvent_* is stronger for the continuum model, because the potential only has a DC component. To quantify this effect, we calculated *SNR_solvent_* as in Equation 2, defining the solvent region by the white portion of the mask in Figure 4C and the protein as the central black region. The results are plotted in Figure 4F where the final *SNR_solvent_* is about a factor of two too strong using the continuum model, while our model closely matches that of experimental data.

### Amplitude contrast

Amplitude contrast can arise from electrons being scattered outside the objective lens aperture, or by removing inelastically scattered electrons using an energy filter. The former is incorporated by applying an aperture function directly to the complex wave function prior to image formation, which results in an attenuation of the expected number of electrons at the detector. This is demonstrated in Figure 5 for a series of aperture diameters and a simulated amorphous specimen with density and thickness as used previously for the “phase plate” with atomic potentials of either carbon (orange circles), phosphorous (grey x’s) or gold (blue squares). The smallest aperture used (0.01 μ) excludes all but the unscattered beam and so is a measure of total transmittance of the simulated layer.

Amplitude losses due to inelastic scattering are incorporated into the multislice formalism via a complex scattering potential, commonly defined as linearly proportional to the real (elastic) potential, for example as in inSilicoTEM. A detailed analysis of why this proportional model is inadequate is found in Dudarev et al. [29]. Our atomically specific inelastic scattering potential is described in the theory section. To demonstrate that it produces the correct amplitude losses, we rearrange Equation 1 to plot the negative-natural logarithm of the expected electron count vs solvent thickness. Fitting this with linear regression gives a readout of the simulated inelastic mean-free path in Figure 6 which closely matches experimental numbers [49].

**Figure 6.**
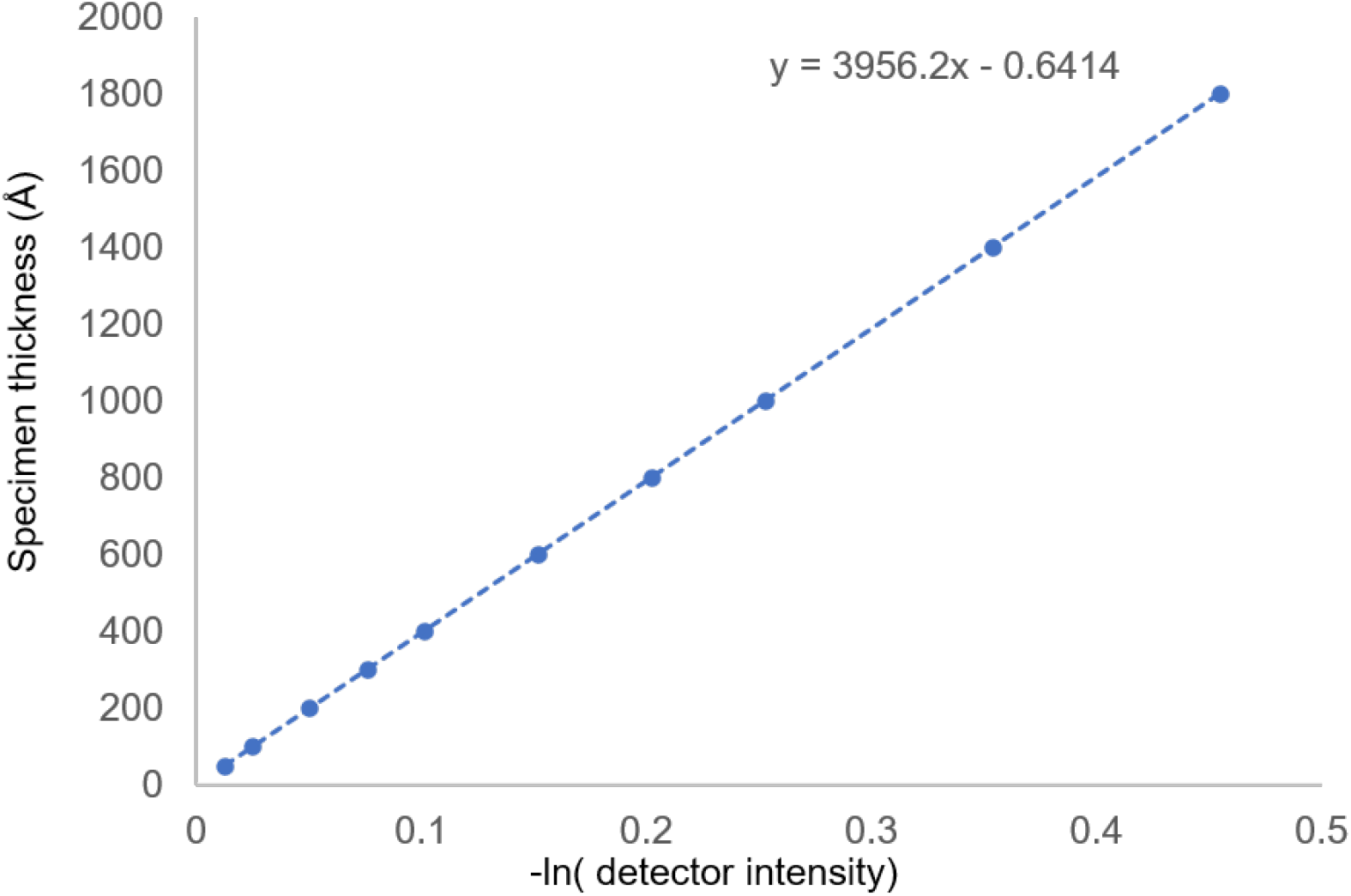
The coarse-grained all-atom solvent model in combination with the inelastic scattering factor for plasmons we derived produce amplitude losses via the complex potential that do not need to be scaled post hoc. The slope is a read out for the inelastic (single-scatter) mean free path in our simulated solvent.

## Discussion

Our simulator implements the most thorough forward model for calculating the interaction between high-energy electrons and radiation-sensitive biological samples demonstrated to date. The improvements described here result from an approximate description of the changes in the specimen due to deposition of energy via inelastic scattering during imaging. This added accuracy in simulating the molecular density produces more realistic image simulations for algorithmic development, but just as importantly, it provides a means to investigate the behavior of complex biological specimens in atomic detail using matched filtering via 2D template matching.

Since the output of the matched filter is sensitive to the spectral distribution of the signal, we can quantify the accuracy of our image formation/damage model by measuring the change in *SNR_mf_*. We found that modeling the water envelop, detector MTF, residual motion blur, and PDB model uncertainty resulted in a higher *SNR_mf_* than could be obtained by optimizing a single B-factor. This analysis is limited by the fact that we cannot strictly disentangle changes to the signal from different envelops that could be mutually compensatory, though this may not be too severe a problem given the differences in the envelopes shown in Figure 3A. A more careful consideration of the impact of different spatial frequencies on *SNR_mf_* may prove useful in addressing this limitation in future work.

Our explicit solvent model, while coarse-grained, allows us to accurately reproduce attenuation due to inelastic losses and amplitude contrast that is spatially variable, and based only on the atomic species and local mass thickness in the simulated specimen. In principle, any configuration of atoms can be simulated by supplying an appropriate PDB file to the simulator. In practice, variable solvent thickness, or other sources of structural noise like regions of hexagonal ice could be included directly into the simulator, however, we leave this for future work. We show a considerable improvement in matching *SNR_solvent_* to experimental data and expect this to improve the ability of models (artificial neural networks especially) trained on simulated data to generalize more readily to experimental data. On visual inspection, the granularity in the solvent appears a bit different than that observed in experiment; we suspect including solvated ions might account for this and plan to include this in future work.

Simulations using an explicit solvent model are computationally very demanding, increasing the number of scattering centers to be considered by a factor of ~200 when simulating single particle image stacks, and up to a factor of 104 when simulating micrographs with well-spaced particles. We have addressed this computational demand via multi-threading in C++. Most of the time taken by the wave-propagation calculation is spent on Fourier transforms. The calculation is currently limited to 4 threads per simulation, independent of the number of threads used in calculating the real-space potentials. To simulate tilted samples, which will have a substantially larger number of slices to propagate, the Fourier transform can become a bottleneck, suggesting a GPU implementation may be beneficial for future work.

## Conclusion

Here we have presented an accurate forward model describing sources of signal attenuation and show how the spectral characteristics of that attenuation improve the output of the matched filter (*SNR_mf_*) as used in template matching for the detection of molecules in cryo-EM images. The *SNR_mf_* is in turn directly related to the mass limit for detection; any improvement in our forward model results in being able to detect smaller particles, which will expand the capacity of template matching in visual proteomics. The increased *SNR_mf_* due to modeling radiation damage is encouraging but should likely be modeled more accurately for the purpose of template matching. We also suggest that our model for inelastic scattering could be improved by direct comparison to experiment using the matched filter. If properly accounted for, we could in principle use this scattering information in particle detection, rather than discarding it using an energy filter.

## Notes

### Competing Interest Statement

The authors have declared no competing interest.

### Summary of Updates

Re-arranged sections in the Introduction and in the Theory sections for clarity based on community feedback. Added detail to some equations and physical constants for easy reference. No significant changes to the Results. Added detail to clarify radiation damage and specimen motion model, related to figure 3.

